# The Mitochondrial Palindrome Scaffold and Architectural Domains

**DOI:** 10.1101/2025.11.11.687892

**Authors:** David A Stumpf

## Abstract

Palindromic motifs comprise 28% of human mitochondrial DNA and form evolutionarily conserved guideposts that organize a persistent structural scaffold. Complementary palindromes of three types delineate mitochondrial architectural domains (MADs) capable of self-organizing dynamic configurations. A library of 423 complementary palindromes generates millions of pairing opportunities across 53,166 human GenBank genomes, revealing a poised system that balances stability and flexibility for constrained change between guideposts. Variations in guidepost flanks distinguish human groups and their homologies to other species, indicating genetic diversity that preceded speciation. These dynamics align with Kauffman’s “poised system” model, in which stability and flexibility coexist to enable evolutionary bursts of diversification. A Neo4j knowledge graph containing 173,431 nodes and 119 million relationships integrated 53,166 human and 2,000 non-human mitochondrial genomes, enabling scalable exploration of mitochondrial architecture. The resulting framework provides a foundation for future studies of mitochondrial evolution, topology, and functional organization.

## Introduction

The mitochondrial phylogenetic tree (phylotree) fails to align with empirical knowledge graph data from full-length mitochondrial sequences (Stumpf 2025a). Rather than reflecting a strictly bifurcating structure, mitochondrial haplotrees reveal a reticulated pattern of descent (Bamshad et al. 2001), with haplotypes clustering in complex and often overlapping ways (Estes 2025). The topology of this graph-like structure remains poorly understood.

Taming of chaos enables life. Holland proposed a fitness model providing adaptability and stability (Holland 2010). Kauffman developed a biopolymer model which self-organized into a *poised system* whose structure maintained a balanced state that is stable enough to resist disruptive changes yet flexible enough to rapidly adjust when conditions shift (Kauffman 1991; 1993).

Multiple lines of evidence from both mitochondrial and ancient nuclear DNA now point toward a reticulated pattern of human evolution with multiple contemporaneous lineages contributed to the genetic makeup of emerging modern humans, rather than a single, unbroken matrilineal lineage. High-coverage ancient mtDNA genomes from diverse geographic contexts reveal deep divergences among early modern human populations, some of which fall outside the variation observed in present-day humans (Peyrégne et al. 2019). Some genetic lineages have deep roots outside of Africa, challenging the out-of-Africa hypothesis (Reich et al. 2010; Sawyer et al. 2012). Early hominoids shared genetic sequences, indicating there was introgression (Villanea et al. 2025) or more distant common ancestors. Unresolved is whether modern humans evolved from a single common ancestor or were genetically diverse as our species emerged.

Native graph databases conforming to the recently published ISO/NEC 39075 GAL Standard (ISO, n.d.) enable precise brute-force analytics and preservation of provenance. The methods outperform other databases when data is highly connected. Using these methods, the current project adds new insights to the themes discussed above. The structuring of data in a graph commonly expose unexpected insights and discoveries which precede hypotheses (Sonawane et al. 2019) (Fecho et al. 2021).

## Results

### Constructing the Knowledge Graph

The knowledge graph (schema in Figure 1) (<u>S1</u>) can be created *de novo* or restore from the Neo4j dump file (<u>S2</u>) and contain 14 node and 21 relationship types with 173,431 nodes, 119 million relationships and their 1,300,498 node and 183,037,885 property values (<u>S3)</u> The 53,166 mt-seq nodes (with Genbank full sequences) and 48,400 human-derived fused_probe nodes (not shown) are legacy items from prior work (Stumpf 2025a). The comparison_seq nodes contain Genbank full sequences of 2,000 non-human individuals from 377 species. Homologies to 677 human-derived fused probes involved 1,003 individuals from 187 species, all mammals (<u>S4</u>)

**Figure 1.**
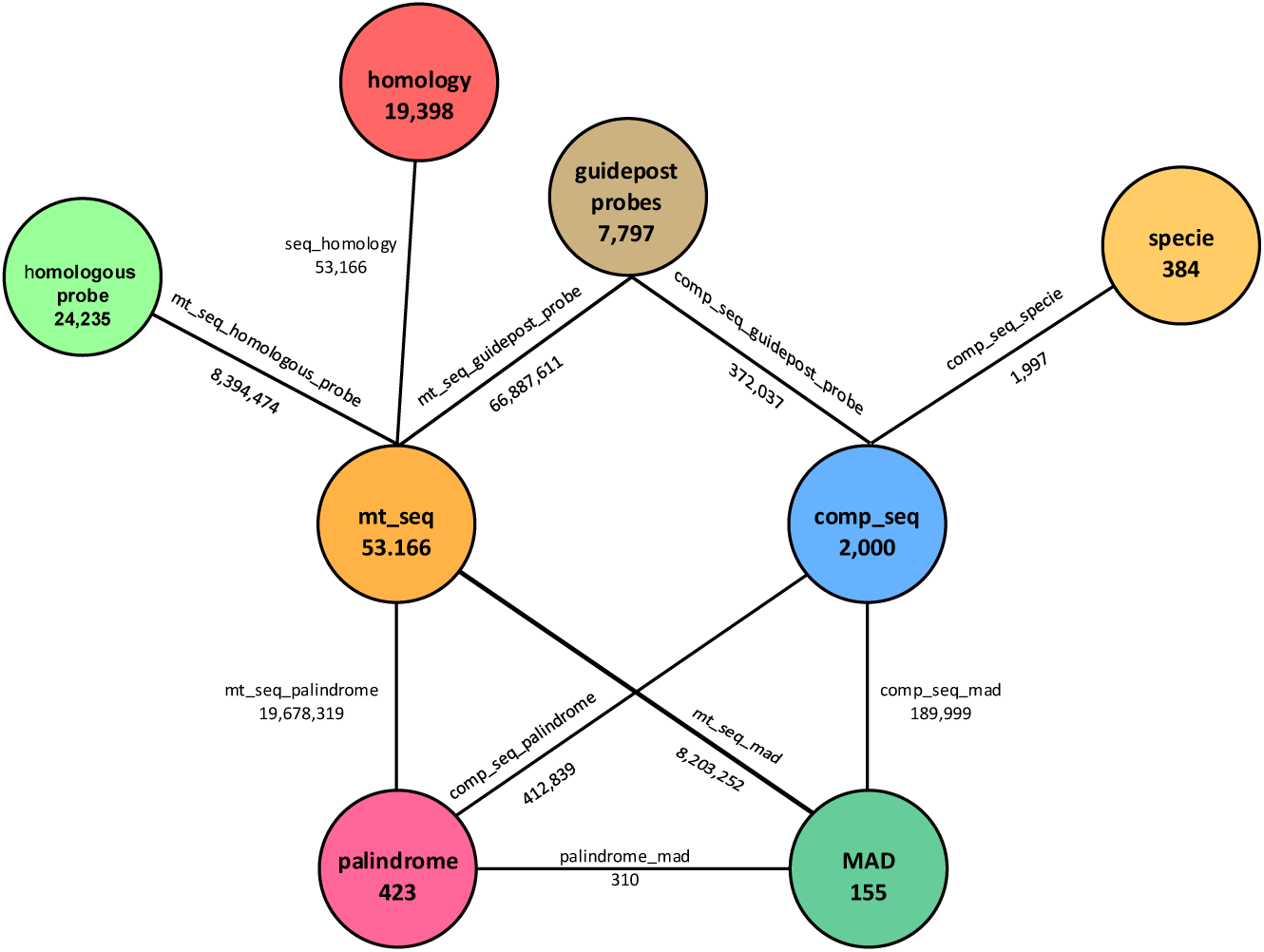
The knowledge graph schema showing key nodes and relationships

The diff-match-patch (DMP) algorithm compared full sequences of human and other species, identifying 8,394,474 seq_homology_probe relationships to 24,235 homologous_probe nodes. Homologies were found to 185 mammalian and 1 fish species (<u>S5</u>). Human sequences (mt_seq) have 2,259,678 relationships to fused probes (not shown), and 66,887,611 guidepost probes.

Palindromes without the flanking nucleotides of fused and homologous probes, were memorialized in 423 nodes, each of which is related to at least one of 410 fused_probe nodes; this is <1% of all fused probes. A palindrome is embedded within 22,629 (93%) of homologous probes (<u>S6</u>) Multiple palindromes were linked by homologies to the comparison_seq and mt_seq nodes through 412,839 compar_seq_palindrome and 19,678,319 mt_seq_palindrome relationships respectively (<u>S3</u>). Palindrome complements were related to each other by 170 palindrom_complement relationships (<u>S7</u>).

### Graph Explorations and Discovery

Palindromes are consistently positioned across Genbank sequences (Table 1). Palindromes exist throughout phylogeny in protozoans, fungi, plants and all phyla of the animal kingdom included in the project (Table 2) (<u>S8</u>). In humans the palindrome bases are 28% of the full sequence (<u>S9</u>). Guidepost_probe sources are 214 (out of the 423) palindromes with a single absolute position in the reference sequence and no flanking bases (flank_size=0). Their 11,086,060 relationships to mt_seq nodes have a property holding the absolute position of the guidepost within the mt_seq. These absolute positions of individual sequence-guidepost pairings are within 5 bases of the guidepost median position in 95% of sequences. In the flank_size=0 guidepost nodes the source and the guidepost probe are the same. Guidepost adjacent flanking bases are retrieved from sequences, creating guidepost probes with 1 to 5 flanking bases on each side of the source palindrome. The unique guidepost probes increase from 214 to 2,392 at flank sizes 0 and 5, respectively. For each guidepost there is a linear relationship between flank size and the observed unique probes, The tight clustering and constrained, rather than random, proliferation of variations suggests a poised system scaffold (<u>S10</u>).

**Table 1.** A snapshot of the palindrome scaffold in 25 Genbank sequences between positions 1000 and 2000. These specific 8 palindromes contain occupy 99.5% of the matrix cells. The “missing” palindrome position has all the palindrome base separated by an insertion

| seq | pos_range | probe_ct | probes |
| --- | --- | --- | --- |
| JQ703707 | 1002-1789 | 8 | [CAGTTGAC, AACACACAA, TACCGCCAT, GATTTAG, AGAGGAGA, CCCAAACCC, ATAGATA, AAGGGAA] |
| JQ704718 | 1002-1789 | 8 | [CAGTTGAC, AACACACAA, TACCGCCAT, GATTTAG, AGAGGAGA, CCCAAACCC, ATAGATA, AAGGGAA] |
| JQ705730 | 1002-1789 | 8 | [CAGTTGAC, AACACACAA, TACCGCCAT, GATTTAG, AGAGGAGA, CCCAAACCC, ATAGATA, AAGGGAA] |
| JX152875 | 1002-1789 | 8 | [CAGTTGAC, AACACACAA, TACCGCCAT, GATTTAG, AGAGGAGA, CCCAAACCC, ATAGATA, AAGGGAA] |
| JX153885 | 1002-1789 | 8 | [CAGTTGAC, AACACACAA, TACCGCCAT, GATTTAG, AGAGGAGA, CCCAAACCC, ATAGATA, AAGGGAA] |
| JX669382 | 1002-1789 | 8 | [CAGTTGAC, AACACACAA, TACCGCCAT, GATTTAG, AGAGGAGA, CCCAAACCC, ATAGATA, AAGGGAA] |
| KC257351 | 1002-1789 | 8 | [CAGTTGAC, AACACACAA, TACCGCCAT, GATTTAG, AGAGGAGA, CCCAAACCC, ATAGATA, AAGGGAA] |
| KC711011 | 1002-1789 | 8 | [CAGTTGAC, AACACACAA, TACCGCCAT, GATTTAG, AGAGGAGA, CCCAAACCC, ATAGATA, AAGGGAA] |
| KF056245 | 1002-1789 | 8 | [CAGTTGAC, AACACACAA, TACCGCCAT, GATTTAG, AGAGGAGA, CCCAAACCC, ATAGATA, AAGGGAA] |
| KF161417 | 1002-1789 | 8 | [CAGTTGAC, AACACACAA, TACCGCCAT, GATTTAG, AGAGGAGA, CCCAAACCC, ATAGATA, AAGGGAA] |
| KF162482 | 1002-1789 | 8 | [CAGTTGAC, AACACACAA, TACCGCCAT, GATTTAG, AGAGGAGA, CCCAAACCC, ATAGATA, AAGGGAA] |
| KF451168 | 1002-1789 | 8 | [CAGTTGAC, AACACACAA, TACCGCCAT, GATTTAG, AGAGGAGA, CCCAAACCC, ATAGATA, AAGGGAA] |
| KF540798 | 1002-1789 | 8 | [CAGTTGAC, AACACACAA, TACCGCCAT, GATTTAG, AGAGGAGA, CCCAAACCC, ATAGATA, AAGGGAA] |
| KJ154638 | 1002-1789 | 8 | [CAGTTGAC, AACACACAA, TACCGCCAT, GATTTAG, AGAGGAGA, CCCAAACCC, ATAGATA, AAGGGAA] |
| KJ669128 | 1002-1789 | 7 | [CAGTTGAC, . . . . ., TACCGCCAT, GATTTAG, AGAGGAGA, CCCAAACCC, ATAGATA, AAGGGAA] |
| KM102018 | 1002-1789 | 8 | [CAGTTGAC, AACACACAA, TACCGCCAT, GATTTAG, AGAGGAGA, CCCAAACCC, ATAGATA, AAGGGAA] |
| KP346044 | 1002-1789 | 8 | [CAGTTGAC, AACACACAA, TACCGCCAT, GATTTAG, AGAGGAGA, CCCAAACCC, ATAGATA, AAGGGAA] |
| KT726047 | 1002-1789 | 8 | [CAGTTGAC, AACACACAA, TACCGCCAT, GATTTAG, AGAGGAGA, CCCAAACCC, ATAGATA, AAGGGAA] |
| KU683166 | 1002-1789 | 8 | [CAGTTGAC, AACACACAA, TACCGCCAT, GATTTAG, AGAGGAGA, CCCAAACCC, ATAGATA, AAGGGAA] |
| KX440324 | 1002-1789 | 8 | [CAGTTGAC, AACACACAA, TACCGCCAT, GATTTAG, AGAGGAGA, CCCAAACCC, ATAGATA, AAGGGAA] |
| KX457382 | 1002-1789 | 8 | [CAGTTGAC, AACACACAA, TACCGCCAT, GATTTAG, AGAGGAGA, CCCAAACCC, ATAGATA, AAGGGAA] |
| KY595624 | 1002-1789 | 8 | [CAGTTGAC, AACACACAA, TACCGCCAT, GATTTAG, AGAGGAGA, CCCAAACCC, ATAGATA, AAGGGAA] |
| KY681004 | 1002-1789 | 8 | [CAGTTGAC, AACACACAA, TACCGCCAT, GATTTAG, AGAGGAGA, CCCAAACCC, ATAGATA, AAGGGAA] |
| MF056020 | 1002-1789 | 8 | [CAGTTGAC, AACACACAA, TACCGCCAT, GATTTAG, AGAGGAGA, CCCAAACCC, ATAGATA, AAGGGAA] |
| MF057058 | 1002-1789 | 8 | [CAGTTGAC, AACACACAA, TACCGCCAT, GATTTAG, AGAGGAGA, CCCAAACCC, ATAGATA, AAGGGAA] |

**Table 2.** Number of unique human palindrome shared across kingdoms.

| kingdom ▼ | Protozoa ▼ | Plantae ▼ | Fungi ▼ | Animalia ▼ |
| --- | --- | --- | --- | --- |
| **Protozoa** | **148** | 122 | 147 | 148 |
| **Plantae** | 122 | **229** | 210 | 229 |
| **Fungi** | 147 | 210 | **293** | 293 |
| **Animalia** | 148 | 229 | 293 | **423** |

Guidepost-probe flank expansion differed among guidepost locations (Figure 2 A-D). Node color intensity corresponds to the magnitude of expansion. The guidepost probe median positions and bases (flask_size=0) shown in the figure are 893: GCCACCG (A); 1548: AGAGGAGA (B); 4723: CATAACCAATAC (C); and 65: GGGGGG (D).

**Figure 2.**
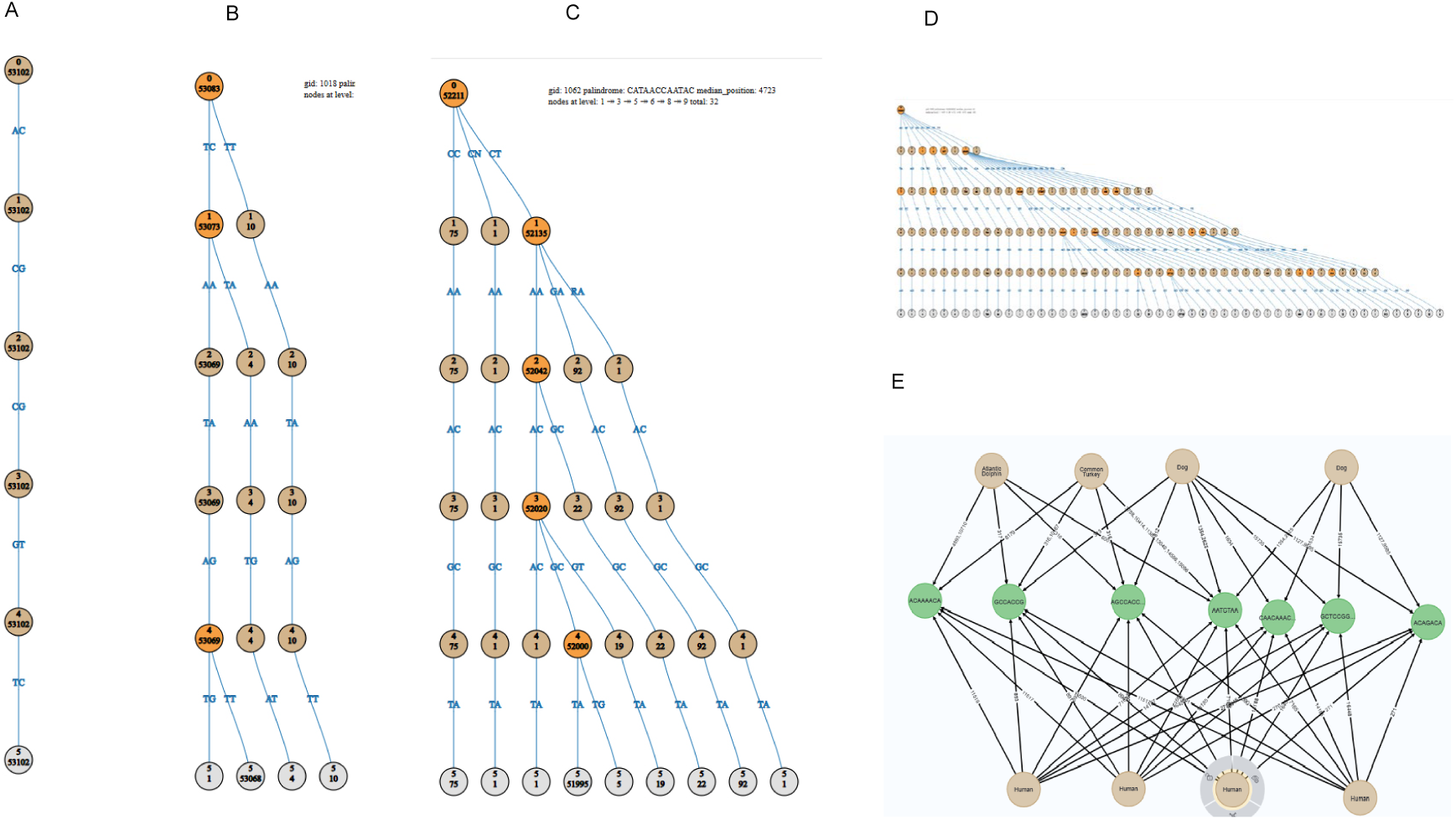
Guidepost-probe flank expansion differs among guidepost locations, ranging from no expansion (A) to limited (B), moderate (C), or major (D) bursts of diversity. Guidepost-probe node labels indicate the number of flanking bases and sequences sharing that probe. Virtual relationships are annotated with the added left and right flanking bases. Nodes are color-coded to highlight bursts. E) Human probes share homologies with other species, including an example of diversity shared among dogs. Individual SVG images are provided in the supplemental materials.

Cross-species comparisons (Figure 2 E) show that human guidepost probes share homologous sequences with other species. Within dogs, multiple homologous probes are present and show diversity among dogs in their homologous sequences. Individual SVG files for each guidepost are provided in the supplemental materials (<u>S11</u>).

Genbank human sequences contain sequences of guidepost probes which are shared with groups of members of other species which form clusters (Figure 3A). The pattern of cluster varies between humans, Neander lands and Denisovans (Figure 3B) (<u>S12</u>).

**Figure 3.**
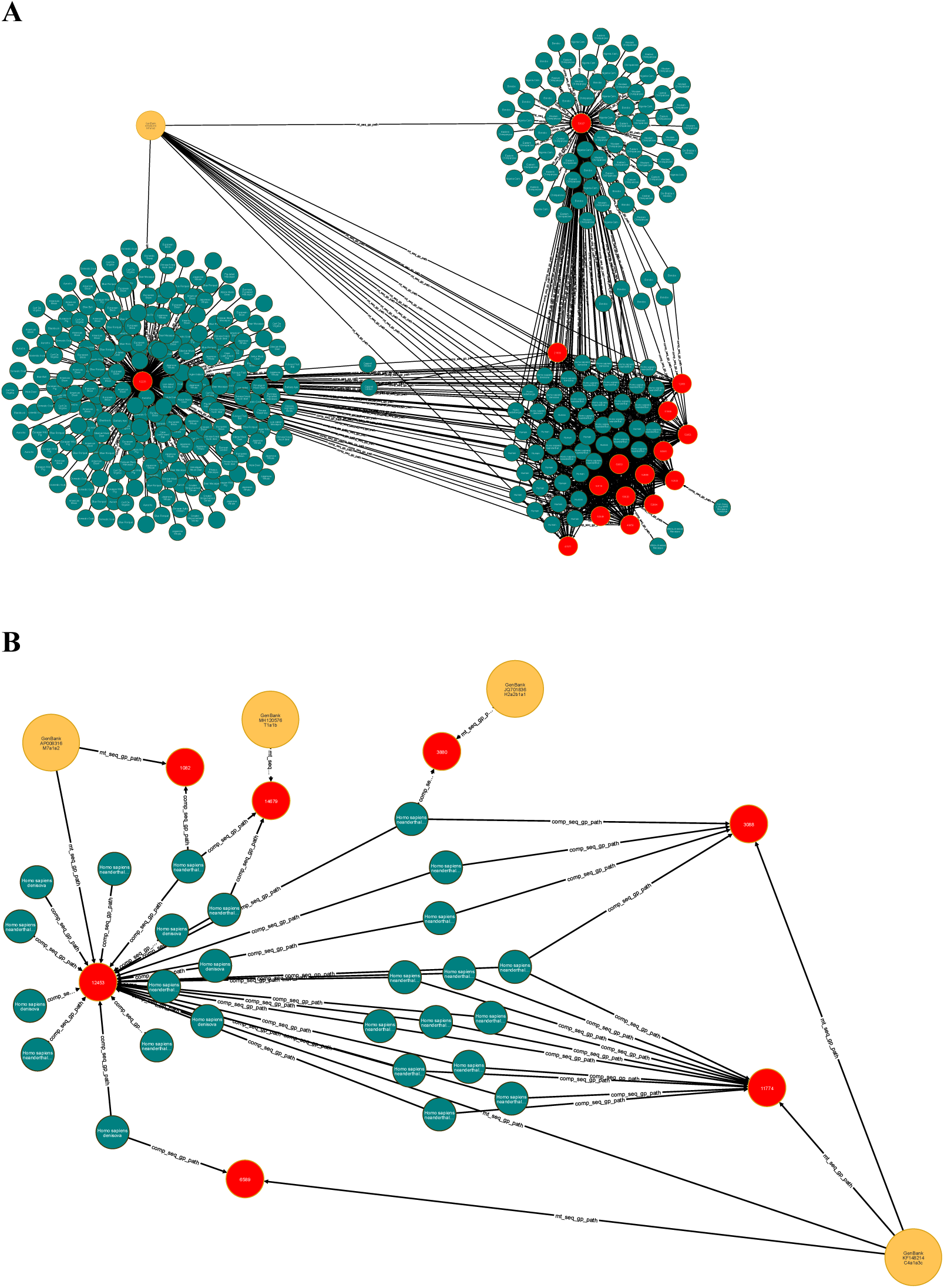
Homologies to other species are not the same for every modern human. A: Human sequences upper eft) has relationships to guidepost_probe nodes around which clusters form. In the upper right are chimpazees and bongos; lower righ early hominoids and lower left a varietyo of other species. This figure will differ between humans. B: Four human nodes connect via guidepost_probe to comparison neanderthal and denosova nodes. Humans connect to different sets of early hominoids via probes shared by the human - hominoid pairs. Human sequence nodes are yellow showing their Genbank name and assigned haplogroup. Guidepost probe nodes are red and contained in the number of sequences noted. Comparison species nodes are green and show the common name of the species.

Three types of palindrome complements were observed within full sequences (see methods); their pairing defined mitochondrial architectural domains (MADs). MAD pairings from humans are present in every kingdom and increase in number in later evolving species (Table 3) (<u>S13</u>). All mammalian orders studied, with the exception of Dasyuromorphia (carnivorous marsupials), have nearly all the 155 MADs seen in humans. Fifty bacteria from 7 evaluated species contain 95% of the palindromes and all of the MADs (<u>S14</u>). MAD nodes were related to the mt_seq and comparison_seq nodes when MAD motif-pairs were both homologous, resulting in 114,846 comp_seq_mad and 4,822,028 mt_seq_mad relationships. These relationships have properties pos1 and pos2 holding lists of positions homologous to motif1 and motif2 respectively.

**Table 3.** Number of distinct human MADs shared across kingdoms.

| kingdom | Protozoa | Plantae | Fungi | Animalia |
| --- | --- | --- | --- | --- |
| Protozoa | **35** | 26 | 35 | 35 |
| Plantae | 26 | **99** | 94 | 99 |
| Fungi | 35 | 94 | **133** | 133 |
| Animalia | 35 | 99 | 133 | **155** |

MADs are consistently positioned in all human sequences. The mean number of palindromes per sequence (154; SD=1) indicates a 99.6% occupancy rate. The MAD pairing type 1, 2 and 3 have 47, 44, 64 nodes and, with overlaps, are in 102, 102, 136 of the MAD pairings (<u>S15</u>). The 154 complements are distributed across the genome (Figure 4) (<u>S16)</u>.

**Figure 4.**
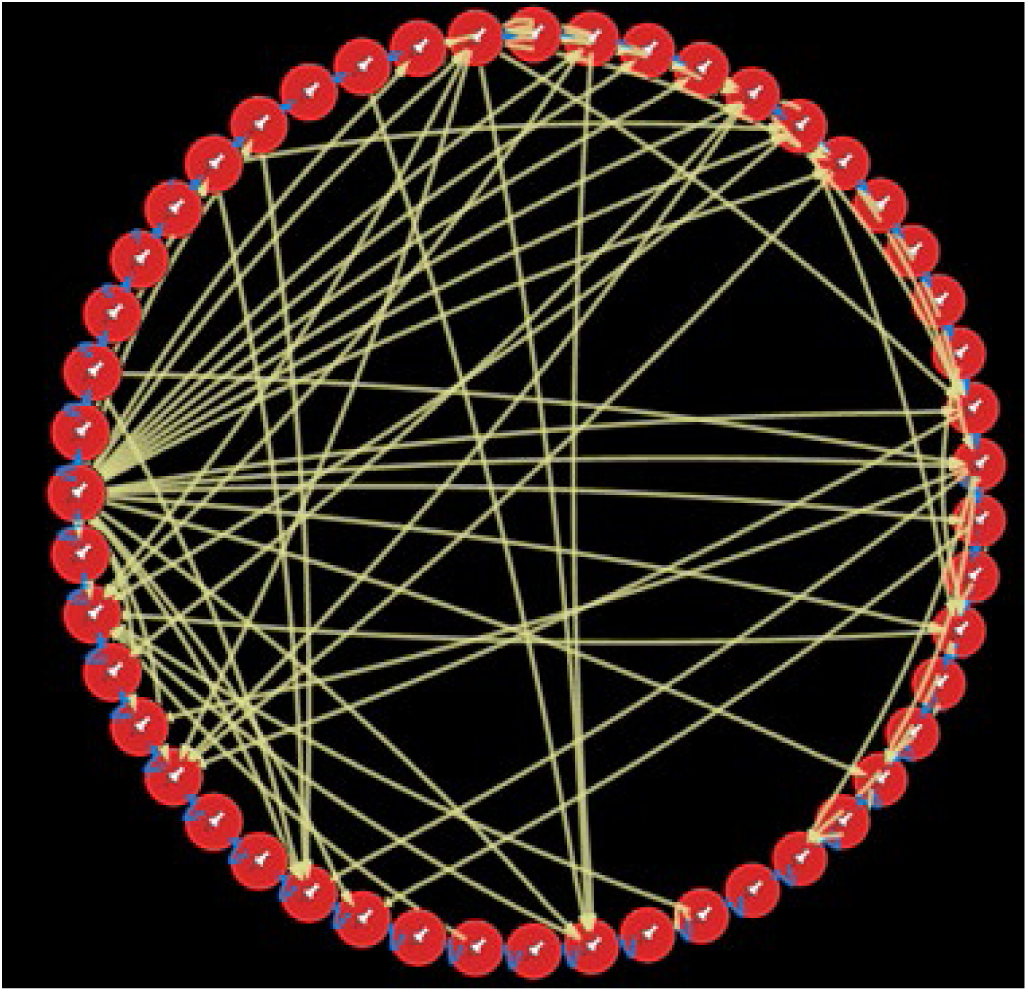
Palindromes (red) linked into a circle by their ordered connections to adjacent palindromes Lines that connect their complements in other position are type 1 (green), 2 (yellow) and 3 (orange). How they associate in vivo is presently unknown.

A subset of guidepost (146 out of 214) became guidepost probes with flank_size=0 and 99% occupancy of all 53,166 Genbank sequences. Designated as “root qualified,” they are most suitable for exploring homologies in other species and identifying root nodes in the phylogeny tree. Pursuing root nodes in an ongoing project. Adding flanking bases to these 146 probes demonstrates diversity before speciation in many species, including humans (<u>S17</u>). These guidepost_probe nodes were co-located with electron transport chain genes and tRNA loci (<u>S18</u>).

The seq_homology relationship, with a 1:1 relationship between mt_seq and homology nodes, creates clusters of human sequences with the same homology json pattern to other species. Cluster size ranges from 1 to 1475 sequences. The sequences in a cluster generally share the same major haplogroup clade. Clades are found in many unrelated clusters; for instance, the H-clade is found in 229 clusters (<u>S19</u>).

### Hypothesis formulation and testing

Observations consistent with a guidepost scaffold led to several hypotheses (Table 3). For each palindrome position, occupancy was tested against p0=0.95 using a one-sample binomial proportion z-test. Across the set, mean occupancy was 0.974 (SD 0.088; n=214; 95% CI 0.962–0.985). A null model of random guidepost positioning was rejected by both a Kolmogorov–Smirnov test and a position-entropy (Shannon) analysis of the sequence-by-position matrix (241 × 53,166; density 0.974) (<u>S20</u>).

Each palindrome anchors a single genomic position. From that anchor, flanking bases extend probes to flank sizes k=0–5. Adding flanks increases discriminatory power: across guideposts, the count of distinct patterns rises ∼linearly with k yet stays far below random base-addition expectations (observed ∼2,600 vs. random 4^10^ = 1,048,576). Linear models fit strongly (high R^2^) for every guidepost, indicating structured, non-random differentiation as k grows. At the per-sequence level, the number of occupied guideposts remains stable across k. Regressions of occupied-count on k produce slopes ≈0 for all sequences; p-values satisfy H_0_: slope =0 across the cohort. The null hypothesis is confirmed. Jaccard-based testing (null: J=1.0; one-sided α=0.005) evaluated within-species diversity in non-human species. Of 377 species, 65 met the sample-size criterion and 48/65 (74%) rejected the null, indicating within-species diversity (<u>S17</u>).

**Table 4.** Hypotheses concerning a scaffold.

|  | Hypothesis |
| --- | --- |
| 1 | Each guidepost is present in nearly all humans. |
| 2 | Guideposts are consistently located at specific positions. |
| 3 | The sequence-guidepost matrix is full. |
| 4 | Guideposts constrain flanking base variation. |
| 5 | Guideposts are persistent across evolutionary time. |

## Discussion

### The Origins of Order

Observed patterns of mitochondrial probe diversity provide quantitative support for Kauffman’s poised-system model of evolution. In *The Origins of Order* (Kauffman 1993), Kauffman used mathematical simulations of binomial networks to show how stable, self-organizing states emerge from structural constraints, generating order without central control. His concept of the “adjacent possible” describes how systems evolve through feasible, incremental innovations while avoiding exploring the entire combinatorial space. Stable mitochondrial guidepost flank expansions linearly increase diversity (slope ≈ 468.7, R² = 0.998) which is constrained as envisioned by Kauffman. Closer inspection reveals faint inflection points, indicating that the apparent linearity arises from the aggregation of many small, stepwise transitions.

The flank expansion options are not random. In graph modeling, the constrained stability has bursts of diversity analogous to Barabási’s formulation of bursty dynamics in complex systems (Barabási 2005) (Goh and Barabási 2008). Eldridge and Gould observed long periods of relative stability interrupted by short, intense intervals of reorganization (Eldredge and Gould 1971). The punctuated evolution, like the present study, produces linear expansion graphs (Venditti and Pagel 2008). However, timelines cannot be inferred from the data in the present study.

### Diversity before Speciation

The palindrome scaffold has ancient origins. Adjacent variations have homologies across species. There are thousands of options for adjacent variations of a specific guidepost and only one is homologous in a pairing of one human and one member of another species. This suggests the variant emerged in a common ancestor of the human and homologous species rather than independently by chance. This mechanism is more likely because other guideposts in a single human have homologies to a variety of different species. Finally, distinct groups of humans share the same pattern of homologies. This evidence indicates humans were diverse as the species emerged in evolution.

Other evidence indicates early hominoids were genetically diverse. Because human mt-DNA has a strict matrilineal inheritance, the diversity of guidepost homologies in a single human cannot be explained by mating between early hominoids. Shared autosomal motifs led to hypothesized matings between early hominoids (Peyrégne et al. 2019). While this is tenable, it may reflect more ancient common ancestors and not interspecies matings.

Diversity before speciation, suggested by mt-DNA analytics, has broad implications for models of evolution. The L7 clade repositioned its members to a new earlier emerging branch (Maier et al. 2022). The present work indicates other clades will be re-rooted and many roots originate before or during human phylogenesis. The classic Linnean phylogeny tree and those developed for population genetics have a single common ancestor for all members of a species. The data suggests otherwise. In contrast to the monoparental mt-DNA inheritance, autosomal DNA can converge to create unique phenotypes of species. Species are also constrained by the ability of members to mate and reproduce.

### Discovery without Hypotheses

Knowledge graph construction connects entities with known relationships fostering exploration. Iterative consolidation of insights in the graph creates positive feedback loops augmenting the graph and preparing it for unanticipated discoveries. Discovery of things hidden in plain sight are common in graph analytics. Often intuitively obvious, discoveries generate testable hypotheses.

The scaffold model followed this path. Fused probes used previously had very few homologies to other species. Direct comparison of human and other species produced probes with homologies between sets of humans and specific mammals. Realizing that most of the homologous probes contained palindromes opened a new line of research and discovering the ubiquitous presence of palindromes and the designation of a subset as guideposts in a scaffold. The model developed was then ready for hypothesis testing.

### The Palindrome Scaffold

Palindromes of 6 to 15 bases, easily identified, appear in one or more positions accounting for 28% of the entire genome. A subset with single locations, remarkably were all at the same position in nearly every human, providing markers or guideposts at discrete positions along traversals of the full sequence. The biological precision of the scaffold and near perfect straight regression lines of variants around the guidepost is extraordinary. These findings mirror the coding principles outlined by Smith-Carpenter et al. (2017), in which complementary nucleoside scaffolds direct synthesis and reversible interactions, foreshadowing the structural information of mitochondrial architecture.

The D-sugar backbone of nucleic acids fixes a single molecular handedness, giving every strand a unidirectional 5′–3′ polarity that enables accurate templating but forbids true molecular mirrors. Palindromic motifs reintroduce mirror symmetry at the informational level: their bases read identically in opposite directions, creating logical reflections within a chemically asymmetric framework. Across 53,166 human mitochondrial genomes, 28 % of bases participate in such palindromic motifs, which form conserved complementary pairs and define a stable yet flexible scaffold for sequence organization. In this way, the chemically imposed directionality of the molecule and the informational symmetry of the sequence act together to shape mitochondrial architecture. This interplay between molecular asymmetry and informational reflection echoes broader theoretical models in which new layers of order arise through the constrained coupling of symmetry and asymmetry in living systems (Malloy et al. 2025; Gleiser et al. 2008). While molecular chirality constrains the chemistry of replication, palindromic pseudo-chirality constrains the organization of information, linking the physical and informational dimensions of the genome.

The RSRS was constructed using statistical (likelihood-based) and rule-based (parsimony) methods applied to existing phylogenetic trees. It captured much of the underlying mitochondrial scaffold and its palindromic loci without explicitly identifying or modeling them. This reliance on statistical inference and parsimony rules perpetuated both their continued use and the interpretive errors they impose on positional accuracy (Li 2014). In contrast, scaffold palindromes provide direct, sequence-anchored targeting of mitochondrial positions through split-alignment methods, achieving nucleotide-level precision unattainable by statistical reconstruction alone (Stumpf 2025a).

### Mitochondrial Architectural Domains

Palindrome pairings with complement motifs within the same sequence form boundaries of architectural domains. An individual pairing could alter the tertiary structure and affect conditions within and adjacent to the domain. Taken together, multiple domain pairings create combinatorial dynamic options, often constrained because they cannot coexist. The complex dynamics within the scaffold consistently exist in each human. The choreography is presently unknown,

Mitochondrial transcription factor A (TFAM) is often described as the principal driver of mitochondrial nucleoid compaction and curvature (Kukat et al. 2015), yet this may invert cause and effect. The preexisting three-dimensional architecture defined by MAD pairings likely predates TFAM evolution and could provide the structural framework into which TFAM binds, stabilizing or constraining rather than creating the configuration.

MAD pairings have analogies to well-known nuclear genome topologically associating domains (TADs) which organize into discrete, highly reproducible structures which constrain genomic folding and gene regulation (Okhovat et al. 2023). Whether MADs behave similarly in nuclear DNA is an open question.

### Precision

The observations reported are very precise. Most biologists intuitively conclude it’s too good to be true; the author agrees. But, in this case the data asserts that it is true. Several factors are involved. The analytics are brute force; there are no statistical best guesses, likelihood computations, or parsimony assumptions. No model is assumed or coercion of data to fit it. A knowledge graph enables unequivocal queries anchored to the provenance of the data. But these factors are trivial in comparison to the underlying truth.

Biological systems achieve a level of precision that at first seems “too good to be true.”” Yet this accuracy is not an artifact of analysis, it reflects an underlying reality. The precision comes from Nature itself. Controlling chaos is the essence of life, and one manifestation of this control is the palindrome scaffold and the architectural stability it provides. Variation is present, but it is dramatically constrained. Changes within the scaffold can enhance fitness, shape fecundity, or contribute to disease.

### Legacy Frameworks

The rCRS, RSRS and Phylotree assembly involved poorly documented curation decisions with little published rationale for the choices made and methods used. Commonly used tools have complex rules buried in code which coerce haplogroup assignments. Remarkably, the RSRS does fully encompass the palindrome scaffold, but tools ignore this by focusing on variations. Clusters based on palindrome patterns usually involve individuals with the same major haplogroup clade, yet they are distributed across unrelated clusters. This reflects differences in the root node(s) used in haplogroup assignments and those found studying homologies to earlier species.

The Modern Synthesis, built on the merger of Mendelian inheritance with Darwinian selection was disrupted by molecular genetics (Rose and Oakley 2007) and non-Mendelian features of human matrilineally inherited mitochondrial disorders (Stumpf 1979). Bacterial and plant phylogenies reveal extensive reticulation, horizontal exchange, and scaffold-like architectural constraints. The ancient origins of the human palindrome scaffold have analogies. The timelines of the Modern Synthesis and its mitochondrial derivatives assume that diversification followed speciation. The ancient origin of the scaffold requires reevaluation of timelines. Evidence also challenges the out of Africa hypothesis and suggests that hominoids have diverse genetic and geographic origins (Reich 2018).

### Opportunities and Feasible Next Steps

This research surfaces new opportunities, suggests testable hypotheses, and provides new tools to expedite exploration. Five are enumerated here.

First, the scaffold reveals missing palindromes in some sequences. Determining whether such losses are neutral, impact fecundity, or cause disease is a natural extension for groups with clinical and reproductive data. Some reported mutations are within palindromes but are insertions rather than the reported point substitutions.

Second, the differing slopes of guidepost flank-size diversity suggest a potential timeline signal, distinct from mutation counts. Calibrating these slopes against demographic and archaeological benchmarks could provide an alternative framework for evolutionary dating.

Third, the scaffold exposes evidence of diversity before speciation. Its conservation across species implies a reticulated, rather than strictly bifurcating, evolutionary process. Extending analyses to early hominins, other animal and plant mitochondria could illuminate how scaffold architecture shaped deep evolutionary history.

Fourth, this work has practical relevance for forensics and genetic genealogy, which maintain the largest collections of human mt-DNA. Scaffold-based methods could enhance identification, classification, and quality control in these applied domains.

Fifth, the scaffold points to a specific class of chemical architectures that may have contributed to life’s origins. Its remarkable stability across kingdoms implies that palindrome-based motifs are not incidental but reflect fundamental organizing principles governing molecular assembly. This perspective aligns with assembly theory (Kempes et al. 2025), which posits that biological systems emerge through the accumulation and selection of reproducible molecular arrangements. In this view, the mitochondrial scaffold exemplifies a deep assembly with an information-rich structure preserved because its architecture supports both stability and adaptive innovation. The pattern echoes Kauffman’s vision of life arising from self-organized molecular order, where architectural constraints channel the balance between persistence and change.

Fifth, the scaffold suggests one specific chemical contributing to the origins of life. Its architectural stability across kingdoms suggests that palindrome-based motifs are not incidental but fundamental organizing principles. This echoes Kauffman’s vision of life emerging from self-organized molecular order where architectural constraints guide stability and innovation. The scaffold therefore may represent one of the earliest molecular frameworks upon which evolution was built.

### Reproducibility Packet

One key product of this research is a set of guidepost palindromes. A python packet with guideposts and full sequences can quickly recapitulate the scaffold and homologies. The packet contains documentation and executes the creation of an SQLite database, loading the data and running several informative queries (Stumpf 2025b). While this does not duplicate all the findings enabled by the knowledge graph, it creates transparency for a broad audience.

### Cautions and Limitations

While the palindrome scaffold model is strongly supported by reproducible analytics and consistent cross-species patterns, several cautions are warranted. The findings rely on publicly available GenBank sequences, which vary in quality and metadata completeness, and thus may incorporate undetected sequencing or annotation errors. The observed precision of palindrome positioning and occupancy could reflect biological constraint but also systematic biases in constructing sequences using reference alignment pipelines. The scaffold and MAD models describe *in silico* structural possibilities rather than demonstrated *in vivo* conformations; biochemical validation will be required to determine whether predicted domain pairings occur within mitochondrial nucleoids. Finally, the knowledge graph approach, though powerful in preserving provenance and enabling brute-force analytics, represents a static snapshot of available data. Future refinements will need to integrate temporal, epigenetic, and transcriptional information to assess how scaffold architecture behaves dynamically within living cells.

## Methods

This study focusses on the topology of a knowledge graph (KG) defined by Genbank full mt-DNA sequences and *in silico* probes derived from them. The foundational methods using human Genbank sequences, described previously (Stumpf 2025a), are expanded to incorporate more probes and full sequences from additional species. Neo4j User Defined Functions (UDFs) create, enhance and analyze the KG and are described and packaged into a downloadable zip file. Executing a single UDF loads the KG by calling sequentially on other functions. Reports render Excel workbooks which reference the UDF, the Cypher queries, and explanations. The datasets used to construct the KG are available in reference files (<u>S21</u>) The Neo4j database can be reconstructed from a dump file (<u>S2</u>) or *de novo* from primary data using a single UDF (mitonet.load;/Load_everything_from_files in <u>S22</u>). The Reproducibility Packet provides validation methods not requiring a graph database (<u>S23</u>). The code was developed in Maven (“Apache Maven,” n.d.) and the pom file contains the required dependencies (<u>S22)</u>.

### Setting up the Neo4j Environment

The knowledge graph was created in a Neo4j Desktop Project v 5.26.1 Enterprise edition (“Download Neo4j Desktop,” n.d.). The APOC package (Neo4j Graph Data Platform, n.d.-a) must be enabled in the desktop and the apoc.config added to the plugin folder. CAUTION: The apoc.config is poorly and often incorrectly documented; use the file in the supplemental materials (<u>S24</u>). and place it in the plugin folder. The Maven compiled jar file is placed in the Neo4j plugin folder. Before compiling, the file folder locations and database name must be configured in the mitonet.neo4jlib.neo4j_info package. If you do these sequentially and verify the database will restart at each step, you can isolate where an issue is encountered. The folder for the import directory is also specified in neo4j_info and by default is a folder withing the dbms stack.

This research makes extensive use of Neo4j User Defined Functions (UDFs) (Neo4j Graph Data Platform, n.d.-e) to connect to, load, enhance, report and manage data provenance. The UDF are coded in java. Each UDF is a java class. Other classes may define data types or other elements instantiated during data processing. The coding was done using Maven, with dependencies in its pom-file and the compiled jar file used in Neo4j by placing it in the plugin folder.

The UDF mitonet.neo4jlib.neo4j_info.neo4j_var() manages the connection to your database and the query UDFs will create sessions which execute query, process return values and returned outputs.

The design is modular. Cypher queries can involve UDFs and UDFs can call other UDFs, including APOC functions. There are brief descriptions of UDFs in Neo4j format and java comments, both of which provide documentation of the UDF.

### Restoring the Neo4j Knowledge Graph from the Dump File

The Knowledge Graph is downloadable (<u>S2</u>). Unzip it and place it in the project folder. This file will then appear at the bottom of the listing in Neo4j Desktop (Neo4j Graph Data Platform, n.d.-d). Click on the dump file and import into your existing DBMS. The restoration will include node, relationships, properties and indices. It requires no reference files and minimal setup. It is the recommended methods for initial work with the graph.

### Loading From Reference Files

A single User Defined Function will load the Knowledge Graph from reference files. The UDF code looks for very specific paths and files. These are set in the mitonet.neo4jLib.neo4jinfo. The author’s code uses an external file to hold the database name, username (default=neo4j), and the password set when the DBMS was created. This can be emulated or these parameters placed directly in neo4j_info. The reference files are unzipped into a folder which is referenced in neo4_info.

The loading is then performed by the UDF mitonet.load.Load_everything_from_files. The load time may be several hours depending on your RAM and processor speed. Tracking messages will be printed out and stored in the import directory. The load process calls on many other UDF or classes.

### Analytics

Strategies are different in knowledge graphs. Initial data, in this case full mt-DNA sequences, including a reference sequence (RSRS) are loaded into the graph. Knowledge Graph (KG) analytic results are memorialized as new nodes, relationships or properties of them. The iterative enhancements meticulously incorporate elements about provenance to make downstream queries more efficient and to enable traversals back to source information.

KG exploration is also iterative and results often enlightening in unanticipated ways. The structure of KG emerges from exploration rather than pre-planning. Discovery occurs without hypotheses. The findings do generate hypotheses which can then be statistically evaluated.

Native graph databases have node, relationship, path and list data types, each with their own set of methods. The relationships in a native graph database can have indexable properties. During graph path traversals across the nodes and relationships properties can be collected into lists or utilized as selection criteria. Neo4j has an APOC plugin (Neo4j Graph Data Platform, n.d.-b) which includes set algebra functions facilitating list analytics including sorting, subtracting (\), finding intersections (∩), checking for specific content (∈), and finding the index of elements within a list (i). The latter is particularly useful in finding position(s) of a probe within a sequence of bases. A KG is generally smaller than relational databases and, in many scenarios, more query-efficient. Efficiency enables brute-force analytics not tenable in other systems.

The KG conforms to the GQL database standard (Enzo 2024). The Cypher query language (Neo4j Graph Data Platform, n.d.-c) in used to build and analyze the KG. The 108 UDF incorporate both Cypher queries and java methods and include 5,795 lines of code.

### Reporting

The UDFs doing the reports are in the packet mitonet.reports. The reports themselves contain the package information at the bottom of the Excel worksheets when generated by a UDF. The Excel reports are formatted with data filters, a split at the top row, auto sized column width and column sums at the bottom. Also, at the bottom are documentation of the UDF package, any Cypher query used and messages to explain the report content and interpretation. The reporting uses an Excel library of UDF code.

The reporting workflow provenance is traceable. Reports are included in the supplemental materials. Supplemental materials are references in the manuscript and links retrieve an Excel report which references the UDF at the bottom. The trace is thus:

Manuscript reference > supplemental materials > report > UDF > UDF documentation

### Statistics

The lists of data in this study were evaluated to determine if they are normally distributed. They usually were not, directing analyses to non-parametric statistics. Graphs were represented in matrices. In the KG 53,166 sequences and 423 palindromes were represented as palindrome present (1) or not (0). The occupancy rate is computed as the portion of each cell to the total. Occupancy of each palindrome across the mtDNA sequence set was tested against the null expectation using a **binomial test** (exact or normal approximation to a one-sample proportion test). In practice, this is equivalent to a z-test for proportions given the large sample size (N > 50,000).

The one-sample Kolmogorov–Smirnov test was used to assess whether positions are uniformly distributes. When the p-value was <0.05 the null hypothesis fails, and the positions are not randomly distributed.

Mutations in DNA involve 4-bases and, if random, would have 4^n^ possibilities. Adding flanks to palindromes exposes variations between full sequences. No single established test directly compares observed versus possible flank patterns. The analysis combines the occupancy (coupon-collector) model, which predicts the mean and variance of distinct patterns if bases were chosen uniformly at random, with a one-sample z-test that standardizes the observed deficit relative to that null. This yields both an effect size (observed-to-possible ratio) and a p-value quantifying the significance of the deviation from the random expectation.

### Reproducibility Packet

The key results can be reproduced in ∼10 minutes using a Python package and data files. The packet deploys an SQLite database rather than the Neo4j graph database used in the study.

## Supporting information

https://arch.library.northwestern.edu/collections/z029p5414?locale=en

## Acknowledgments

The Mitomap team provided haplogroup assignments for most sequences used. Richard Haas, Teepu Siddique and W. Wesley Johnston provided feedback on the manuscript. Steven J Coker, Robert Marvin and Alan Wilson tested the Reproducibility Packet. The author acknowledges the use of OpenAI’s ChatGPT v 5 for assistance in preparing this manuscript and some of the code. This required detailed prompts using the author’s domain knowledge to focus the chats.

## Data Availability

All materials required to reproduce this work, including datasets, analysis scripts, and derived files, are available at the Northwestern University Library ARCH server (“eLife Supplemental Materials,” n.d.). Each dataset has an assigned DOI to ensure permanent, citable access. In addition, contextual links within the manuscript provide direct access to the relevant data and code *in situ*.

