## Supplementary material for "The Mitochondrial Palindrome Scaffold and Architectural Domains": https://arch.library.northwestern.edu/collections/z029p5414?locale=en

Supplemental Materials

David A Stumpf, MD, PhD

The Supplemental Materials are available at the Northwestern University ARCH Server from which they can be download and in some cases unzip and password protected (password is Woodstock!Mitonet).

Hot links to these material are in the manuscript and labeled a S# where the numbering is sequential.
